# Whole genome sequencing identifies a novel factor required for secretory granule maturation in *Tetrahymena thermophila*

**DOI:** 10.1101/042085

**Authors:** Cassandra Kontur, Santosh Kumar, Xun Lan, Jonathan K. Pritchard, Aaron P. Turkewitz

**Author notes:** made equal contributions.

## Abstract

Unbiased genetic approaches have a unique ability to identify novel genes associated with specific biological pathways. Thanks to next generation sequencing, forward genetic strategies can be expanded into a wider range of model organisms. The formation of secretory granules, called mucocysts, in the ciliate *Tetrahymena thermophila* relies in part on ancestral lysosomal sorting machinery but is also likely to involve novel factors. In prior work, multiple strains with defect in mucocyst biogenesis were generated by nitrosoguanidine mutagenesis, and characterized using genetic and cell biological approaches, but the genetic lesions themselves were unknown. Here, we show that analyzing one such mutant by whole genome sequencing reveals a novel factor in mucocyst formation. Strain UC620 has both morphological and biochemical defects in mucocyst maturation, a process analogous to dense core granule maturation in animals. Illumina sequencing of a pool of UC620 F2 clones identified a missense mutation in a novel gene called *MMA1* (*M*ucocyst *ma*turation). The defects in UC620 were rescued by expression of a wildtype copy of *MMA1*, and disruption of *MMA1* in an otherwise wildtype strain generated a phenocopy of UC620. The product of *MMA1*, characterized as a CFP-tagged copy, encodes a large soluble cytosolic protein. A small fraction of Mma1p-CFP is pelletable, which may reflect association with endosomes. The gene has no identifiable homologs except in other Tetrahymena species, and therefore represents an evolutionarily recent innovation that is required for granule maturation.

## Background

All eukaryotes possess a network of membrane-bound organelles that underlie wide-ranging cellular activities. While the basic features of this network are conserved, there is phylogenetic evidence that specific pathways have tended to experience high levels of innovation over evolutionary time, including pathways directly involved in protein secretion(1-4). From a cell biological perspective, such innovations are interesting because they may underlie specialized secretory responses. Dense core granules (DCGs) in animal cells are secretory organelles that are adapted for the storage of bioactive peptides(5, 6). These peptides can subsequently be released in response to extracellular stimuli, a phenomenon called regulated exocytosis(7). Studies of DCG biogenesis, particularly in mammalian endocrine cells, have detailed a biosynthetic pathway in which newly formed DCGs undergo extensive maturation before they are competent for cargo exocytosis(8, 9). Maturation involves pathway-specific machinery. For example, key proteases dedicated to generating bioactive peptides, the prohormone convertases, were evolutionarily derived from a trans-Golgi network protease called KEX2/furin that is conserved within the Opisthokont lineage (including both fungi and animals) but not demonstrably present in other eukaryotic lineages(9, 10). More recently, genetic dissection of DCG biogenesis in invertebrates is uncovering numerous additional factors. Some of these are conserved within metazoa, without any clearly homologous genes in other lineages(11, 12). Thus, mechanisms underlying the formation of dense core granules in animals are likely to depend in part on lineage-restricted innovation.

Ciliates are a large clade of unicellular eukaryotes, many of which are striking in their behavioral and structural complexity. This complexity includes a remarkable array of secretory vesicles that contribute to functions including predation, predator deterrence, and encystation(13). The ciliate vesicles show marked structural and functional similarities to metazoan dense core granules(14). However, as members of the SAR (Stramenopile/Alveolate/Rhizaria) lineage, Ciliates are very distantly related from Opisthokonts(15). This immense evolutionary divergence raises the possibility that dense core granules in Ciliates could have arisen largely independently from those in animals(16, 17).

Biochemical and other molecular data for ciliate granules is largely limited to two species belonging to the Oligohymenophorean branch, *Tetrahymena thermophila* and *Paramecium tetraurelia*(18-21). The ciliate granule cargo proteins, like many proteins in endocrine granules, are acidic, bind calcium with low affinity, and form large aggregates within the secretory pathway, suggesting that compartment-specific aggregation may be a widespread mechanism for sorting to secretory granules(22-26). Moreover, the ciliate proteins subsequently undergo proteolytic maturation via endo-and exoproteolytic processing, similar to cargo proteins in metazoan endocrine granules(24, 27). However, neither the ciliate cargo proteins nor the processing enzymes are homologous to their functional counterparts in metazoans, suggesting that unrelated proteins evolved to produce similar functions within the secretory pathway(28-31). That idea is also consistent with identification of a set of genes required in *P. tetraurelia* for granule exocytosis, which are lineage-restricted rather than conserved(32-35). Similarly, while *T. thermophila* and *P. tetraurelia* express large families of classical eukaryotic trafficking determinants such as Rab GTPases and SNAREs, they lack any clear homologs of some specific subtypes that are associated with secretory granules in metazoans(36-38). Taken together, the current data are consistent with the idea that similar functions in granule formation can be provided in ciliates and animals by paralogs that arose independently within the same gene families, or by unrelated genes.

One potentially powerful approach to expanding our catalog of genes involved in ciliate granulogenesis is forward genetics, which has provided a key tool in dissecting mechanisms of membrane trafficking in budding yeast and other organisms(39, 40). Importantly, screens based on random mutagenesis are free of the bias inherent in candidate gene approaches. In *T. thermophila*, mutants with a variety of defects in secretory granule biogenesis have been generated using nitrosoguanidine, and characterized using genetic and cell biological approaches, but none of the underlying genetic lesions has been identified(41-44). However, advances in high throughput sequencing should make it possible to identify the causative mutations via whole genome comparisons, as has recently been demonstrated for a *T. thermophila* mutant in ciliary basal body orientation(45).

The secretory granules in *T. thermophila*, called mucocysts, are filled primarily with proteins of the Grl (Granule lattice) family(46). The Grl proteins undergo proteolytic processing that is required for morphological maturation of newly synthesized mucocysts(24, 47-49). Mucocysts also contain a second family of abundant proteins(31, 50). The best-studied member is Grt1p (Granule tip), whose name derives from the polarized distribution of this protein in mature mucocysts(43). Both Grl proprotein processing and Grt1p polarization are largely blocked in an exocytosis-deficient mutant, generated by nitrosoguanidine mutagenesis, called UC620(43). The UC620 mutation showed Mendelian inheritance expected for a recessive allele, and the mucocyst maturation defects were partially suppressed under starvation conditions(43). We have now used whole genome sequencing, applied to F2 progeny of UC620, to identify the genetic lesion in this mutant. Analysis of the gene, which we call *MMA1* for *M*ucocyst *Ma*turation 1, illustrates the power of forward genetics to uncover ciliate-restricted innovations required for secretory granule biogenesis.

## Materials and Methods

### Cells and Cell Culture

*Tetrahymena thermophila* strains used in this work are listed in Table 1. Unless otherwise stated, cells were cultured in SPP (1% proteose peptone, 0.2% dextrose, 0.1% yeast extract, 0.003% sequestrene (ferric ethylenediaminetetraacetic acid) and starved in 10mM Tris buffer, pH 7.4, both at 30°C while shaking at 99 rpm. PP for cells grown in 96 well plates also included 250μg/ml penicillin G, 250μg/ml streptomycin sulfate, and 0.25μg/ml amphotericin B fungizone (Gibco). For most uses, cells were grown overnight to medium density (1.5-3×10^5^ cells/ml) in a volume of SPP equal to one fifth of the nominal culture flask volume. Cell densities were determined using a Z1 Beckman Coulter Counter. Cells in 96-well or drop plates, including cells under drug selection, were grown for 3 days at 30°C in moisture chambers. Scoring and screening of cells was done on an inverted microscope at 100x magnification.

**Table 1:** Strains used in the work

| Strain | Drug Resistance<br>[micronuclear genotype<br>(macronuclear<br>phenotype)] | Exocytosis<br>Phenotype | Mating Type | Notes |
| --- | --- | --- | --- | --- |
| B2086 | *mpr1-1/mpr1-1* (mp-s) | exo<sup>+</sup> | II | parental |
| CU428 | *mpr1-1/mpr1-1* (mp-s) | exo<sup>+</sup> | VII | parental<br>(mutagenized) |
| CU427 | *chx1-1/chx1-1* (cy-s) | exo<sup>+</sup> | VI | initial outcross |
| CU438 | *pmr1-1/pmr1-1* (pm-s) | exo<sup>+</sup> | IV | outcross, this paper |
| IA264 | *gal1-1/gal1-1* (dg-s) | exo<sup>+</sup> | II | outcross, this paper |
| UC620 | *chx1-1/chx1-1; mpr1-1/mpr1-1* (cy-r, mp-r) | exo<sup>-</sup> |  |  |
| F1 | CHX/*chx1-1*; MPR/*mpr1-1*; GAL/*gal1-1*, exo<sup>+</sup>/exo<sup>-</sup> (cy-r, mp-r, dg-r) | exo<sup>+</sup> |  |  |
| CU428<br>*MMA1*-CFP | *mpr1-1/mpr1-1* (mp-s); (cy-r) | exo<sup>+</sup> | VII | *MMA1*-CFP inducibly expressed in CU428 background |
| UC620<br>*MMA1*-CFP | *chx1-1/chx1-1; mpr1-1/mpr1-1* (cy-r, mp-r); (cy-r) | exo<sup>+</sup> |  | *MMA1*-CFP inducibly expressed in UC620 background |
| $\Delta mma1$ | *mpr1-1/mpr1-1* (mp-s); (pmr-r) | exo<sup>-</sup> | VII | *MMA1* disruption in CU428 background |

### Genetic Crosses

Cultures of strains to be mated were grown to 1.5-3 × 10^5^ cells/ml, washed twice, and resuspended in starvation medium, 10mM Tris, pH 7.4, to a final density of 2 x 10^5^ cells/ml. Unless otherwise specified, cells were pelleted in 50ml conical tubes at ~600-1000g for 1 minute. 10 ml of the cells were starved overnight (12-18 hours) at 30°C in 100mm Petri dishes to initiate sexual reactivity. Equal volumes (5ml) of cells to be mated were then gently mixed, within the 30°C incubator, in a new Petri dish. Mating cells were incubated at 30°C for 5-8 hours in a moist chamber, and then refed with an equal volume of 2%PP (2% proteose peptone, 0.003% sequestrene) to generate karyonides(51-53).

### Selecting Progeny

Beginning one hour after refeeding, single mating pairs were isolated by mouth pipetting into individual 30ul drops of SPP in a Petri dish containing 48 drops in an 8 x 6 grid array. To maximize the number and diversity of progeny established, exconjugants were also isolated by distributing 50ul/well of mating pairs into a 96 well plate containing 50μl/well SPP(54). Cells were grown, then replicated into separate 96-well plates containing 100μl/well 2% PP supplemented with the appropriate drug to eliminate drug sensitive parentals. Selection with cycloheximide (chx) and 6-methyl purine (6-mp) were done at 15μg/ml (both from 1000x stock; chx stock in 100% methanol), paromomycin (pms) at 100μg/ml (from 1000x stock), and 2-deoxygalactose (2-dgal) at 2.5mg/ml (from 50x stock)(54). After 3 days growth, plates were scored for drug resistance to identify progeny. Cells selected with 2-dgal were scored after 7 days.

### Obtaining drug-sensitive F1 assortants

In outcrosses where one parent contributed an allele conferring drug resistance (chx1-1), exconjugant progeny were initially heterozygous in both the MIC and MAC. Through phenotypic assortment, some subsequent vegetative progeny will lose all copies of the resistance allele in the MAC. We identified these progeny by serial passaging combined with drug sensitivity tests, as described in (53).

### Mating Type Testing

Bacterized medium was made by adding *Pseudomonas syringae* to a 25ml flask of SPP, and growing overnight at 30°C with shaking at 225 rpm. This culture was diluted 50 fold in sterile water to constitute 2% bacterized peptone (BP). Clones to be tested were replicated into a 96 well plate with 50μl/well 2% BP and grown for 3 days. A 100μl inoculum of each tester strain (MT II, III, IV, V, VI, or VII), taken from an overnight culture, was added to 10ml 2% BP in Petri dishes and grown for 3 days. Tester strains were obtained from the Tetrahymena Stock Center (https://tetrahymena.vet.cornell.edu/). At that time, 50μl/well of the tester strains were separately added to each unknown and plates were incubated at 30°C. A control matrix of each tester separately mated to each of the others and itself was also included. Between 4-8 hours after mixing, cells were scored for pairing. The mating type was defined for clones that paired with 5 of the 6 mating type testers, as equivalent to that of the sole tester strain with which it failed to pair.

### Testing assortants for exocytosis competence using Alcian Blue

Cells were grown in SPP, replicated into a 96 well plate with 50μl/well 2% BP, and grown for 3 days. Capsule formation was induced by adding 50μl/well 0.02% Alcian blue in 0.5mM CaCl_2_solution to each well, followed by the immediate addition of 25μl of 2% PP. Wells were scored for the presence of Alcian blue-stained capsules. Clones showing no capsule formation whatsoever were then retested in bulk culture, before being selected for genomic DNA extraction. For this, cells were grown to 3x10^5^ cell/ml in 25ml SPP, washed twice and resuspended in 10mM Tris, pH 7.4, then starved overnight (12-18 hours) at 30°C. Cells were concentrated to 3ml and transferred to a new 125ml flask, stimulated by rapid addition via syringe of 1ml 0.1% Alcian blue, then diluted after 15 sec with 45ml 0.25% PP + 0.5mM CaCl_2_. Cells were washed once in 50ml Tris, concentrated to 2ml, and resuspended in 25ml Tris for screening by microscopy.

### Genomic DNA Extraction

Cells were grown to 3x10^5^cells/ml in 25 ml SPP, and starved for 18-24 hours in 10mM Tris-HCl, pH 7.5. 1.5ml of cells in an Eppendorf tube were concentrated by pelleting to 50μl. 700μl urea buffer (42% w/v urea, 0.35M NaCl, 0.01M Tris pH 7.4, 0.01M EDTA, 1% SDS) was added followed by gentle shaking, and then adding 0.1mg/ml Proteinase K for a 5 min incubation at 50°C. 750μl phenol:chloroform:isoamyl alcohol (25:24:1) was added, and the tube contents mixed by inversion and then spun for 15 min at 3500rpm. The top, aqueous layers were transferred to new tubes using pipette tips with ends cut off, and the extraction repeated. The top layers were then extracted with an equal volume of chloroform:isoamyl alcohol (24:1), and mixed with a one-third volume of 5M NaCl. DNA was precipitated with an equal volume of isopropyl alcohol, gently spooled onto a hooked glass pipette and transferred to a new non-stick Eppendorf tube (Eppendorf LoBind Microcentrifuge tube^®^). DNA was washed and pelleted twice with 1ml 70% ethanol, and the final pellet left to air dry for 5-10 min. DNA was resuspended in 25μl 1x TE, pH 8.0, treated with 2μl RNAse A (10mg/ml, Fermentas) overnight at 55°C, and stored at −20°C.

#### Disruption of MMA1 to generate Δmma1 strains

*MMA1* (TTHERM_00566910) was replaced in the Macronucleus with the neo4 drug resistance cassette, generously provided by K. Mochizuki (IMPA, Vienna, Austria) via homologous recombination with the linearized vector pUC620MACKO-neo4. We amplified ~600 bp of the upstream and downstream flanks of *MMA1* using the primer pairs 034 and 035, and 036 and 037, respectively (Supplementary Table S1), and cloned them into the SacI and XhoI sites of the neo4 cassette respectively, using In-Fusion cloning kit^®^ (Clontech, Mountain View, CA). CU428 cells were then biolistically transformed with the final construct pUC620MACKO-neo4, linearized with NotI and SalI(25, 28). Transformants were selected on the basis of paromomycin resistance, then serially transferred for 3-4 weeks in increasing drug concentrations to drive fixation of the null allele.

### RT-PCR Confirmation of *MMA1* Disruption

Cultures were grown to 1.5-3.0x10^5^ cells/ml, washed, and starved for 2h in 10mM Tris pH 7.4. Total RNA was isolated as per manufacturer’s instructions using RNeasy Mini Kit (Qiagen, Valencia, CA). The presence of *MMA1* transcripts was assayed with the OneStep RT-PCR kit (Qiagen) using primers (083 and 073, Supplemental Table S1) to amplify ~650bp of the *MMA1* gene. Gene knockout was confirmed by the continued absence of the corresponding transcripts after ~2 weeks of growth in the absence of drug selection (4-5 serial transfers/week). To confirm that equal amounts of cDNA were being amplified, control RT-PCR with primers specific for the αTubulin gene were run in parallel.

### Vector construction and expression of the *MMA1*-CFP gene fusion

The *MMA1* gene was cloned into the pBSICC Gateway vector, a gift from D. Chalker (Washington University, St. Louis, MO), with primers listed in Supplementary Figure S1. Briefly, *MMA1*, minus the stop codon, was PCR amplified using primers 091, 092, and the product first cloned into the entry vector pENTR/D-TOPO (Invitrogen, Grand Island, NY). CACC was added to each forward primer in order to allow directional cloning into pENTR-D. Subsequently, LR Clonase II (Life Technologies) was used to recombine the *MMA1* coding sequence into a destination vector (pIBCC) containing an *MTT1*-inducible CFP tagged expression cassette cloned upstream of a cycloheximide-resistant rpl29 allele. For transformation, the construct was linearized with BaeI and SpeI and biolistically transformed into the wildtype CU428 or mutant UC620 cell lines(28).

### Biolistic Transformation

*Tetrahymena* were transformed by biolistic transformation as previously described (25, 28). Afterward, filters were transferred to a flask with 50ml prewarmed (30°C) SPP without drug and incubated at 30°C with shaking for 4 hours. Transformants were then selected by adding paromomycin (120 μg/ml with 1μg/ml CdCl_2_), or cycloheximide (12μg/ml). Cells were scored for drug resistance after 3 (paromomycin) or 5 (cycloheximide) days. Transformants were serially transferred 5 days a week in decreasing concentrations of CdCl_2_ and increasing concentrations of drug. In all cases, selection was for at least 2 weeks before further testing(55).

#### Immunofluorescence

To visualize mucocysts, cells were fixed, permeabilized with detergent, immunolabeled with mAb 5E9 (10%) or mAb 4D11 (20%) hybridoma supernatant, and analyzed as described previously (28, 56). After 2h induction by 2μg/ml CdCl_2_, Mma1p-CFP fusion protein was visualized using Rabbit anti GFP (Invitrogen) (1:400), respectively, followed by Alexa 488-conjugated anti-Rabbit antibody (1:250). For simultaneous imaging of Mma1p-CFP and Grl3p, cells were costained using the mAb 5E9 as previously described and with the polyclonal anti-GFP antibody (Life Technologies) diluted 1:400 in 1%BSA. Cells were then coincubated with the 2° antibodies Texas red-coupled goat anti-mouse IgG diluted 1:99 and 488-coupled donkey anti-rabbit IgG diluted 1:250 in 1%BSA. Cells were imaged using a Leica SP5 II Confocal Microscope and image data were analyzed as previously described(28). Images were captured with the LAS_AF confocal software (Leica) for Windows 7. Image data were colored and adjusted for brightness/contrast using ImageJ.

#### Dibucaine Stimulation

Dibucaine stimulation of exocytosis was performed as described previously (57).

#### Subcellular fractionation

Cells were grown to 3x10^5^/ml and then transferred into 10mM Tris, pH 7.4. Transgene expression was induced using 0.25μg/ml CdCl_2_ for 2h at 30°C. Cells were chilled and centrifuged (1k x g) in a clinical centrifuge for 1min. All subsequent steps were at 4°C. Cells were resuspended and washed once in Buffer A (20mM HEPES-KOH pH 7.0, 38mM KCl, 2mM MgCl_2_ and 2mM EGTA) and the pellet volume measured. The pellet was resuspended in three volume of Buffer B (20mM HEPES-KOH pH 7.0, 38mM KCl, 2mM MgCl_2_ and 2mM EGTA, 0.3M Sucrose) containing protease inhibitor cocktail tablet (Roche). Cells were passed through a ball-bearing cell cracker with nominal clearance of 0.0004 inches. The homogenate was centrifuged for 30 min at 10,000g. To separate cytosolic and membrane fractions, that supernatant was further centrifuged for 1h at 100,000g. After centrifugation, supernatant (cytosolic) and pellet (membrane) fractions were dissolved in SDS-PAGE buffer and incubated for 15 min at 90°C.

### Tricholoracetic acid precipitation of whole cell lysates

~3×10^5^ cells were pelleted, washed twice with 10mM Tris pH 7.4, and precipitated with 10% trichloroacetic acid (TCA). Precipitates were incubated on ice for 30min, centrifuged (18k x g, 10min, 4°), washed with ice-cold acetone, re-pelleted (18k x g, 5min, 4°) and then dissolved in 2.5x SDS-PAGE sample buffer.

### Immunoprecipitation

CFP-tagged fusion protein was immunoprecipitated (for western blot) from detergent lysates using polyclonal rabbit anti-GFP antiserum as described previously(58).

### Western blotting

Samples were resolved by SDS-PAGE and transferred to 0.45μm PVDF membranes (Thermo Scientific, Rockford, IL). Blots were blocked and probed as previously described (59). The rabbit anti-Grl1p, rabbit anti-Grl3p, rabbit anti polyG (60) and mouse monoclonal anti-GFP (Covance, Princeton, New Jersey) 1° antibodies were diluted 1:2000, 1:800,1:10,000 and 1:5000 respectively. Protein was visualized with either ECL Horseradish Peroxidase linked anti-rabbit (NA934) or anti-mouse (NA931) (Amersham Biosciences, Buckinghamshire, England) 2° antibody diluted 1:20,000 and SuperSignal^®^ West Femto Maximum Sensitivity Substrate (Thermo Scientific, Rockford, IL).

#### Gene expression Profiles

Expression profiles were derived from the *Tetrahymena* Functional Genomics Database (http://tfgd.ihb.ac.cn/), with each profile normalized to that gene’s maximum expression level(61, 62).

#### In silico analyses

Alignment of protein sequences was performed using CLUSTALX (1.8) with default parameters.

### Library construction

DNA was sonicated to produce a 300-400 bp distribution, and then enzymatically end-repaired and A-tailed using Illumina’s standard TruSeq DNA library preparation. These products were ligated to Illumina Truseq adapters. Libraries, following indexing via PCR, were size selected to remove adapter and PCR dimers, pooled, and co-sequenced on two lanes of the Illumina HiSeq 2500 using a 2 × 100 base pair format. The unique sequences incorporated during library construction were subsequently used to identify the source of each read, using postsequencing bioinformatics.

#### Whole genome sequencing

Sequencing of macronuclear DNA library was performed using Illumina HiSeq 2500 by the University of Chicago Genomics Facility at the Knapp Center for Biomedical Discovery (KCBD). The sequencing process followed the manufacturer’s instructions, and the sequence files (fastq) were produced using the Illumina demultiplexing software CASAVA (v1.8). A total number of 269 million paired-end reads (2 x 101 bp) were generated for 5 different strains of *Tetrahymena thermophila*. The F2 lines with the mutation of interest were sequenced to ~70 fold genome coverage. The parental strain, i.e. the wildtype background upon which the mutations were initially induced, was sequenced to ~256 fold genome coverage. The three wild type strains that were used to generate the F2 lines were sequenced to ~65, 73, and 58 fold genome coverage respectively. Genome sequencing of *Tetrahymena* primarily reflects the Macronuclear genome, in which genes are generally present at ~45 copies compared to the 2 copies in the Micronucleus. Because the Micronucleus represents the germline nucleus, only Micronuclear alleles are transmitted to progeny(63). However, because all parental lines are wildtype regarding exocytosis except for UC620 itself, we made the simplifying assumption that bulk DNA sequencing of mutant vs. parental lines would permit us to identify the causative mutation in UC620.

#### Sequence alignment

Reads from total genomic DNA sequencing were mapped to *Tetrahymena thermophila* Macronuclear genome sequence released by Broad institute using the Burrows-Wheeler Aligner software (BWA) version 0.5.9(64-66). Default parameters were used when running bwa aln except the following: 1). “Maximum edit distance” was set to “2”; 2). “Maximum number of gap opens” was set to “0”; 3). “Number of threads” was set to “8”; and 4). “Iterative search” was disabled. The output files were converted to bam files using the “sampe” utility of the BWA software with default parameters and the SAMtools software version 0.1.18(67). SAMtools was then used to sort the bam files and remove PCR duplicates. Picard Tools version 1.92 was used to add group names to the bam files with parameter “VALIDATION_STRINGENCY” set to “LENIENT” (http://picard.sourceforge.net). The resulting bam files were indexed by SAMtools.

#### Variants discovery

The Genome Analysis Toolkit (GATK) version 2.5-2 was applied to identify variants with total genomic DNA sequencing data of the five samples(68). The following steps were taken in this procedure. 1). Realignment of reads using RealignerTargetCreator and IndelRealigner tool of the GATK package to correct the misalignment caused by site mutations, insertions and deletions. 2). Variants were called using the HaplotypeCaller tool of the GATK package with the “out_mode” set to “EMIT_ALL_CONFIDENT_SITES”, “stand_call_conf” set to “50.0” and “stand_emit_conf” set to “30.0”. A total number of 67,256 variants were called with this procedure. Among these, 23,493 were Single-nucleotide variants (SNVs), which result in a SNV density of ~0.23 per kilo base pairs for a genome of a mappable size of 103,014,375bp(65).

#### Candidate screening

The candidate causal variants of the cell exocytosis dysfunction phenotype were selected with the following steps. 1). Variants with missing values for the genotypes were filtered. 2). Candidate sites had to be homozygous in all strains, and the mutant F2 strains’ genotype must be different from the genotype of the parental strain and wild type strains. A total number of 28 such candidate sites were found. Among these 28 variants, 10 were on chromosome 1, 7 were on chromosome 2, 4 were on chromosome 3, 1 was on chromosome 4, 4 were on chromosome 5 and 2 were undefined.

Data and reagents from this work will be made freely available upon request for non-commercial purposes.

## Results

### Generation of F2 clones for sequencing

The concentration of nitrosoguanidine used for mutagenesis to generate UC620 is expected to produce a large number of mutations(43). For that reason, we could not usefully compare the genomes of UC620 to that of the unmutagenized parent strain, as this would be confounded by the large background of single nucleotide variants (SNVs) unrelated to the mutation of interest. To partially overcome this problem, we outcrossed UC620 with strains bearing useful drug resistance markers, and used drug resistance to select progeny that would be heterozygous in the germline Micronuclei for the UC620 mutation. We then mated several progeny clones with one another, and derived the F2 progeny from isolated mating pairs. Since the UC620 mutation is recessive, ¼ of the progeny, namely those homozygous for the mutant alleles, should display the UC620 phenotype. We therefore focused on matings producing the expected 1:3 ratio of mutant:wildtype exocytosis phenotypes among the progeny.

Identifying the homozygous mutant F2 progeny was complicated by the phenomenon of phenotypic assortment, which derives from the amitotic division of the Macronucleus during cytokinesis(63). In particular, even within clones of cells heterozygous for the UC620 mutation, some cells will lose most or all wildtype alleles in the Macronucleus, due to the random assortment of Macronuclear alleles at each cell division(69). For this reason, we judged the homozygous mutant clones to be those showing no wildtype phenotypes whatsoever, in tests performed initially in 96-well plates and confirmed in bulk cultures. We obtained 25 such clones, derived from 2 matings.

DNA was prepared from the individual clones, and then pooled for sequencing. In parallel, we prepared and sequenced DNA from the strains used during the initial mutagenesis to create UC620, as well as from the strains used for the outcrosses described above.

### A Tetrahymenid-restricted gene on chromosome 1 represents a candidate for the mutated gene in UC620

A total of 28 SNVs were identified that were homozygous in the mutant pool but not present in the parental strains. The criteria used in SNV discovery are detailed in Materials and Methods. Of the 28, 10 fell on chromosome 1, to which the UC620 mutation had previously been mapped. Four of these chromosome 1 candidates fell within previously annotated genes. Among these, we focused on TTHERM 00566910 based on its expression profile. An online database of *T. thermophila* gene expression allows one to visualize the expression profiles of all genes in this organism, over a range of culture conditions(61). Using this database, we previously discerned that a large number of genes associated with mucocyst biogenesis had nearly identical expression profiles(28, 70). TTHERM 00566910, which we have called *MMA1* for *M*ucocyst *ma*turation, is expressed at less than twice the corrected background for the whole genome dataset(62). However, the shape of the expression profile was strikingly similar to that of known mucocyst-associated genes (Figure 1A). This profile was not shared by other genes harboring SNVs on chromosome 1 (not shown). On this basis, *MMA1* emerged as the prime candidate for the gene underlying the defect in UC620.

**Figure 1.**
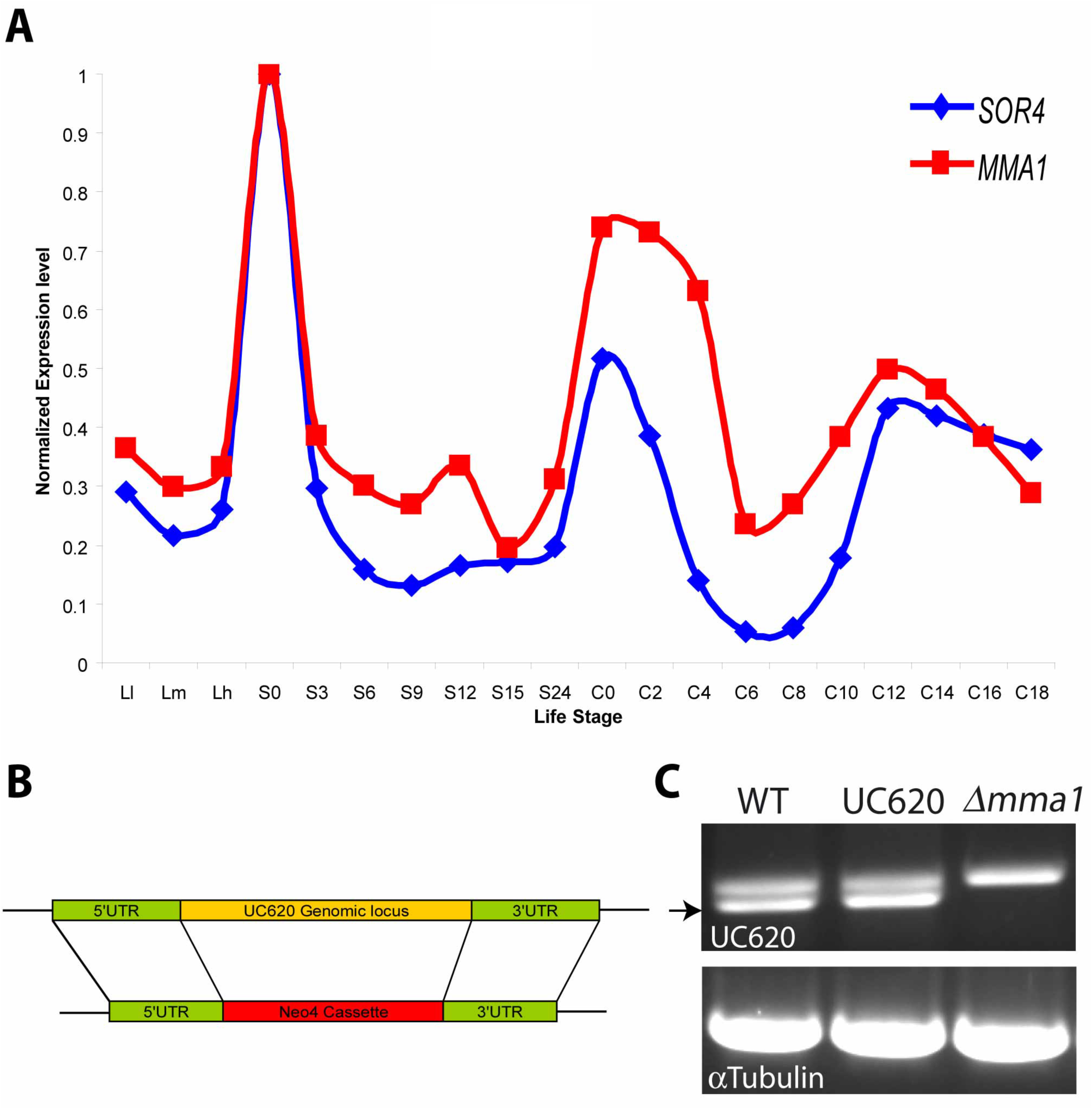
Expression profiling and genomic knockout of *MMA1*. (A) The expression profile of *MMA1* (red trace) is highly similar to that of *SOR4* (blue trace), which encodes a receptor required for mucocyst biogenesis. The profiles of transcript abundance under a variety of culture conditions, derived via hybridization of stage-specific cDNAs to whole genome microarrays, are from the Tetrahymena Functional Genomics Database (http://tfgd.ihb.ac.cn/). In the plots shown, each trace was normalized to that gene’s maximum expression level. The culture conditions sampled at successive time points represent growing (L-l, L-m, and L-h), starved (S-0, S-3, S-6, S-9, S-12, S-15, and S-24), and conjugating (C-0, C-2, C-4, C-6, C-8, C-10, C-12, C-14, C-16, and C-18) cultures. For details on the sampling conditions, see (62). (B) Schematic of *MMA1* Macronuclear gene knockout construct. Replacement of the Macronuclear *MMA1* gene by the Neo4 drug resistance cassette was targeted by homologous recombination. (C) Confirmation of *MMA1* Macronuclear knockout by RT-PCR. RNA was extracted from WT, UC620, and *Δmma1*, converted to cDNA, and PCR amplified using *MMA1*-specific primers listed in Supplemental Table S1. A 1% ethidium bromide-stained agarose gel is shown. The *MMA1* product (indicated by arrow), which was confirmed by sequencing, was present in WT and UC620 samples but absent in *Δmma1*. Parallel amplification of the αTubulin gene from all samples was used to control for sample loading.

*MMA1* has clear homologs in several recently sequenced Tetrahymena species (*T. malaccensis*, *T. ellioti*, and *T. borealis*) (Supplementary figure 2). Alignment of these genes, together with extensive RNAseq data from *T. thermophila*, facilitated confident assignment of exon-intron boundaries, as well as start and stop codons, for this previously undescribed gene. Based on this annotation, the SNV identified in UC620 changed the *MMA1* stop codon (TGA) to TGT, which encodes cysteine. The mutation therefore potentially results in an aberrant polypeptide product. Except for the homologs in these other Tetrahymenids, we could not identify *MMA1* homologs in any other species.

### Disruption of *MMA1* blocks synthesis of docked mucocysts

To ask whether a defect in *MMA1* might account for the mutant phenotype in UC620, we disrupted *MMA1* in wildtype cells by homologous recombination with a drug resistance cassette (Figure 1B). The resulting *Δmma1* cells had no detectible *MMA1* transcript (Figure 1C). They grew at wildtype rates, indicating that the gene is not essential for normal growth under laboratory conditions.

Strikingly, the *Δmma1* cells demonstrated a strong defect in mucocyst exocytosis. The complete failure to release mucocyst contents upon cell stimulation was identical to that of UC620, when tested with the secretagogues Alcian blue (not shown) or dibucaine (Figure 2A). This defect was due to the failure of *Δmma1* cells to synthesize mature docked mucocysts, as revealed by indirect immunofluorescence using antibodies against two mucocyst cargo proteins, Grl3p and Grt1p. The *Δmma1* cells in growing cultures did not accumulate Grl3p in docked mucocysts (Figure 2B). Instead, as in UC620, the Grl3p localized to relatively homogeneous cytoplasmic vesicles.

**Figure 2.**
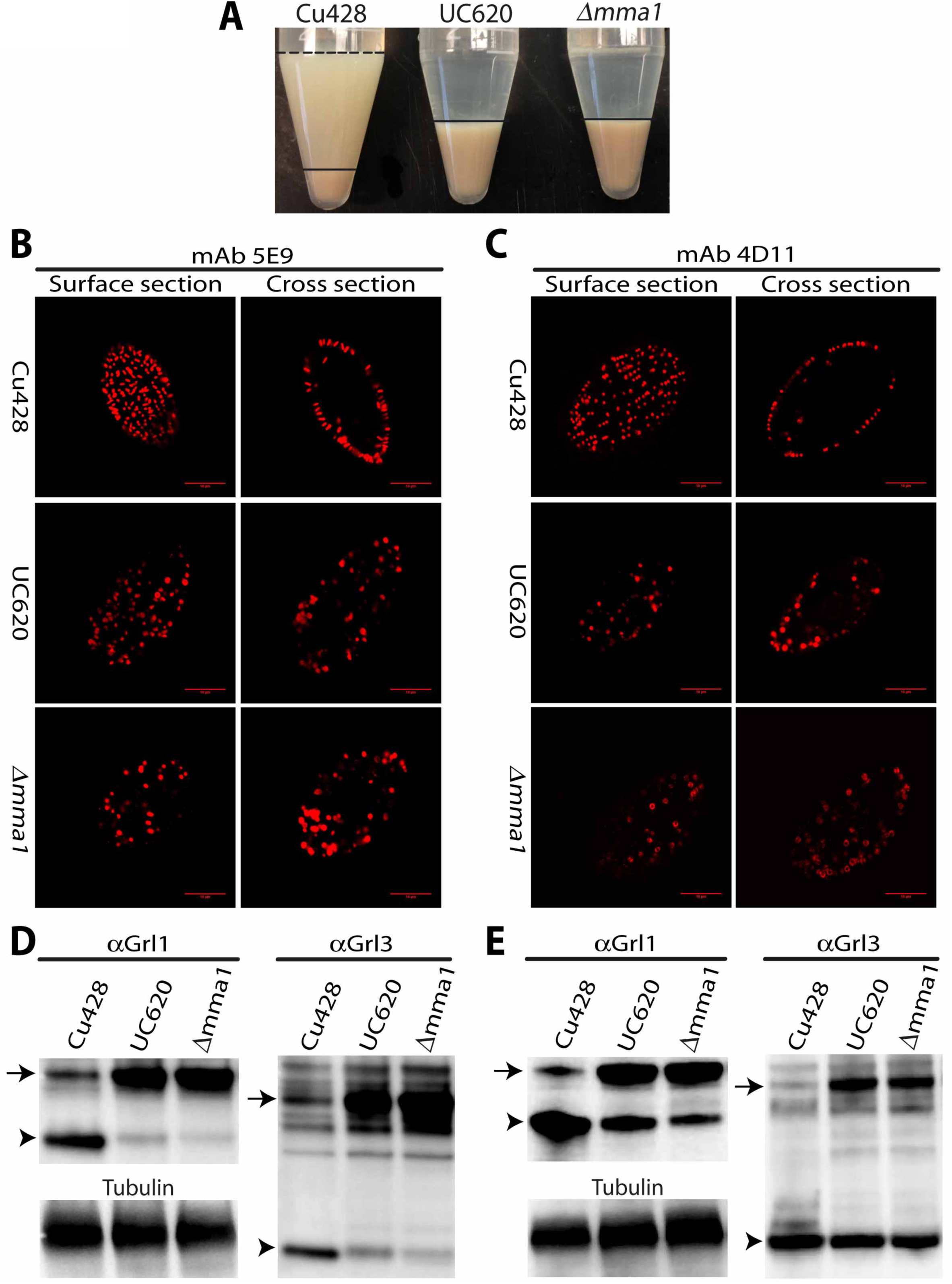
*MMA1* knockout produces a phenocopy of the UC620 mutation. (A) Equal numbers of CU428 wildtype, UC620 and *Δmma1* cells were stimulated with dibucaine, which promotes global mucocyst exocytosis, and then centrifuged. In wildtype cells, this results in a pellet of cells with an overlying flocculent consisting of the released mucocyst contents. For clarity, the flocculent layer is delineated with a dashed line at the upper border, and an unbroken line at the lower border. In contrast, neither UC620 nor *Δmma1* produces detectible flocculent. The pellet in wildtype (layer beneath solid line) is smaller than in the mutants because many wildtype cells are trapped in the flocculent. (B-C) Wildtype, UC620 and *Δmma1* were fixed, permeabilized, and stained with antibodies against two mucocyst core proteins, Grl3p (left panels, 5E9 antibody) and Grt1p (right panels, 4D11 antibody). Shown are optical sections of individual cells, at the cell surface and a cross section. Wildtype CU428 accumulate docked mucocysts, visible as elongated vesicles that are highly concentrated at the cell periphery. UC620 and *Δmma1*, in contrast, do not form elongated vesicles, and Grl3p and Grt1p are found in vesicles throughout the cells. The Grt1p signals in both UC620 and *Δmma1* often appear localized to the vesicle perimeter, while in wildtype cells the signal is localized to the docked mucocyst tips. The scale bars represent 10 μm. (D-E) Whole cell lysates of growing (panel D) or 6 hr starved (panel E) cells were resolved by 4-20% SDS-PAGE, and transfers were immunoblotted with antibodies against Grl1p or Grl3p. The unprocessed (pro-Grl) and processed forms of the Grl proteins are indicated by arrows and arrowheads, respectively. In CU428, Grl proteins accumulate primarily in the fully processed form. In contrast, the pro-Grl forms predominate in growing cultures of UC620 and *Δmma1*. For both UC620 and *Δmma1*, the defect in processing proGrl proteins is partially rescued under starvation conditions (panel E). To demonstrate equivalent loading, all samples were immunoblotted in parallel with anti-tubulin antibody (panels D,E). Each lane in panels D and E represents 10^3^ cell equivalents.

Similarly, a 2^nd^ mucocyst protein, Grt1p, was mislocalized in both UC620 and *Δmma1* cells. In wildtype cells, Grt1p resides in docked mucocysts (Figure 2C). In both UC620 and *Δmma1*, Grt1p instead localizes in cytoplasmic vesicles, frequently appearing as a ring around the vesicle periphery (Figure 2C).

### *Δmma1* cells are defective in proteolytic maturation of mucocyst contents

In UC620, the mislocalization of Grl3p and other Grl-family proteins is accompanied by their aberrant biochemical maturation: the Grl pro-protein precursors largely fail to undergo proteolytic processing that occurs during mucocyst maturation in wildtype cells(43). The *Δmma1* cells showed the same defect in pro-Grl processing, shown for two different Grl proteins (Figure 2D). Moreover, the processing defect in *Δmma1* was partially suppressed when cells were transferred to starvation conditions, as we had previously reported for UC620 (Fig 2E)(43). These results are consistent with the idea that *MMA1* represents the affected gene in the UC620 mutant.

### Mma1p appears to be cytosolic and partially membrane-associated

A hydropathy plot based on the primary sequence of Mma1p did not reveal either a signal sequence, as would be expected for a protein translocated into the secretory pathway, nor any hydrophobic stretches that could function as transmembrane helices (not shown). Thus the protein is likely to be cytosolic. We expressed a CFP-tagged copy of *MMA1* under the control of the strong and inducible *MTT1* (metallothionein 1) promoter(71). We resorted to this over-expression strategy because the very low level expression of wildtype *MMA1* made it impossible to detect the expression of 3xGFP-tagged *MMA1* expressed at the endogenous locus (not shown). Mma1p-CFP, immunoprecipitated from whole cell lysates, appeared by Western blotting as a band of the expected size (Figure 3A). We fractionated cells expressing Mma1p-CFP by cracking them with a ball bearing homogenizer, and then subjecting the cleared lysate to high-speed (100k x g) centrifugation. The protein was primarily found in the soluble fraction, but roughly 10% was found in the high-speed pellet (Figure 3B). Visualization of the CFP in these cells by indirect immunofluorescence showed a large number of small puncta present throughout the cell, but no significant co-localization with mature docked mucocysts (figure 3C). Taken together, these results suggest that Mma1p-CFP is a cytosolic protein that is partially associated with small vesicles.

**Figure 3.**
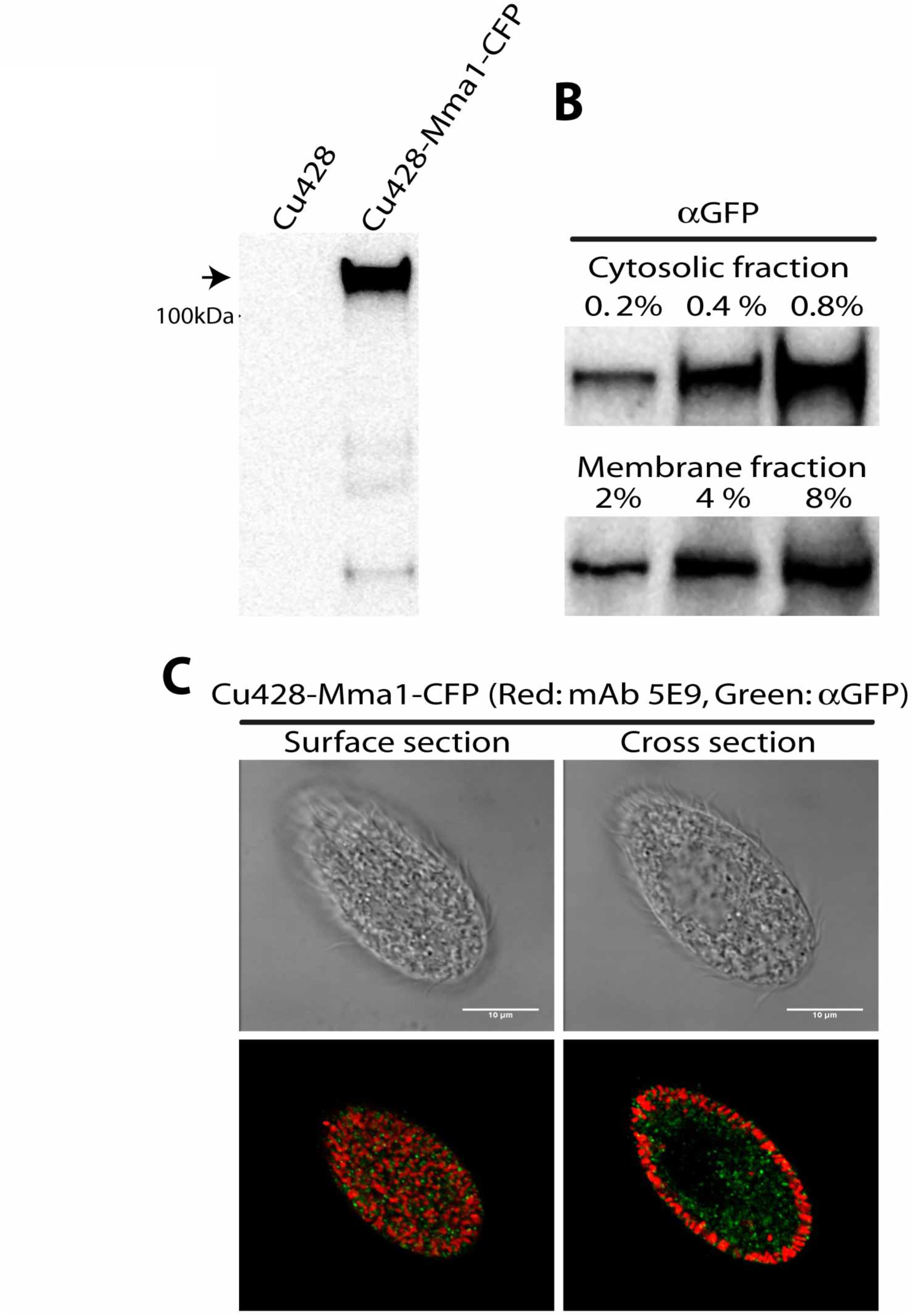
Expression and characterization of Mma1p-CFP. (A) Expression of *MMA1*-CFP. Mma1-CFP fusion protein was immunoprecipitated from detergent cell lysates using anti-GFP antiserum, with wildtype CU428 cells processed in parallel. Immunoprecipitates were subjected to 4-20% SDS-PAGE, and PVDF transfers blotted with anti-GFP mAb. An immunoreactive band (arrowhead) of the size expected for the Mma1p-CFP fusion is seen only in cells expressing this construct. (B) Cells expressing Mma1p-CFP were grown to 3×10^5^/ml, and then fractionated into soluble (cytosolic) and pelletable (membrane) fractions, as described in Materials and Methods. Samples were separated by SDS-PAGE and immunoblotted with anti-GFP mAb. Three different loadings are shown for each sample, corresponding to 0.2-0.8% of the entire cytosolic fraction, and 2-8% (i.e., 10X higher cell equivalents) of the membrane fraction. Approximately 10% of the Mma1p-CFP is found in the membrane fraction. (C) After induction as in panel A, cells were fixed, permeabilized, and immunolabeled with rabbit anti-GFP and mAb 5E9, followed by 2° fluorophore-coupled Abs. The scale bars represent 10μm. Mma1p-CFP (green) appears in small puncta through the cytoplasm, with no significant overlap with docked mucocysts (red). Wildtype cells processed in parallel showed no significant labeling in the green channel (not shown).

### Expression of Mma1p-GFP rescues the UC620 mutant

To ask whether the CFP-tagged copy of Mma1p was active, we introduced the identical construct in UC620 cells. Since the mutation in UC620 behaves as a recessive allele, the defects in these cells should be rescued by the expression of a wildtype copy of the affected gene. A polypeptide of the expected size for the fusion protein was detected by Western blotting, and was localized to a large number of small cytoplasmic puncta (Figure 4A, B). Importantly, UC620 cells expressing *MMA1*-GFP were indistinguishable from wildtype in both mucocyst accumulation and in pro-Grl processing (Figure 4C, D, E). The full rescue of UC620 cells by expression of *MMA1* provides confirmatory evidence that whole genome sequencing has allowed us to identify the genetic lesion in UC620, and that *MMA1* represents a novel factor required for mucocyst maturation.

**Figure 4.**
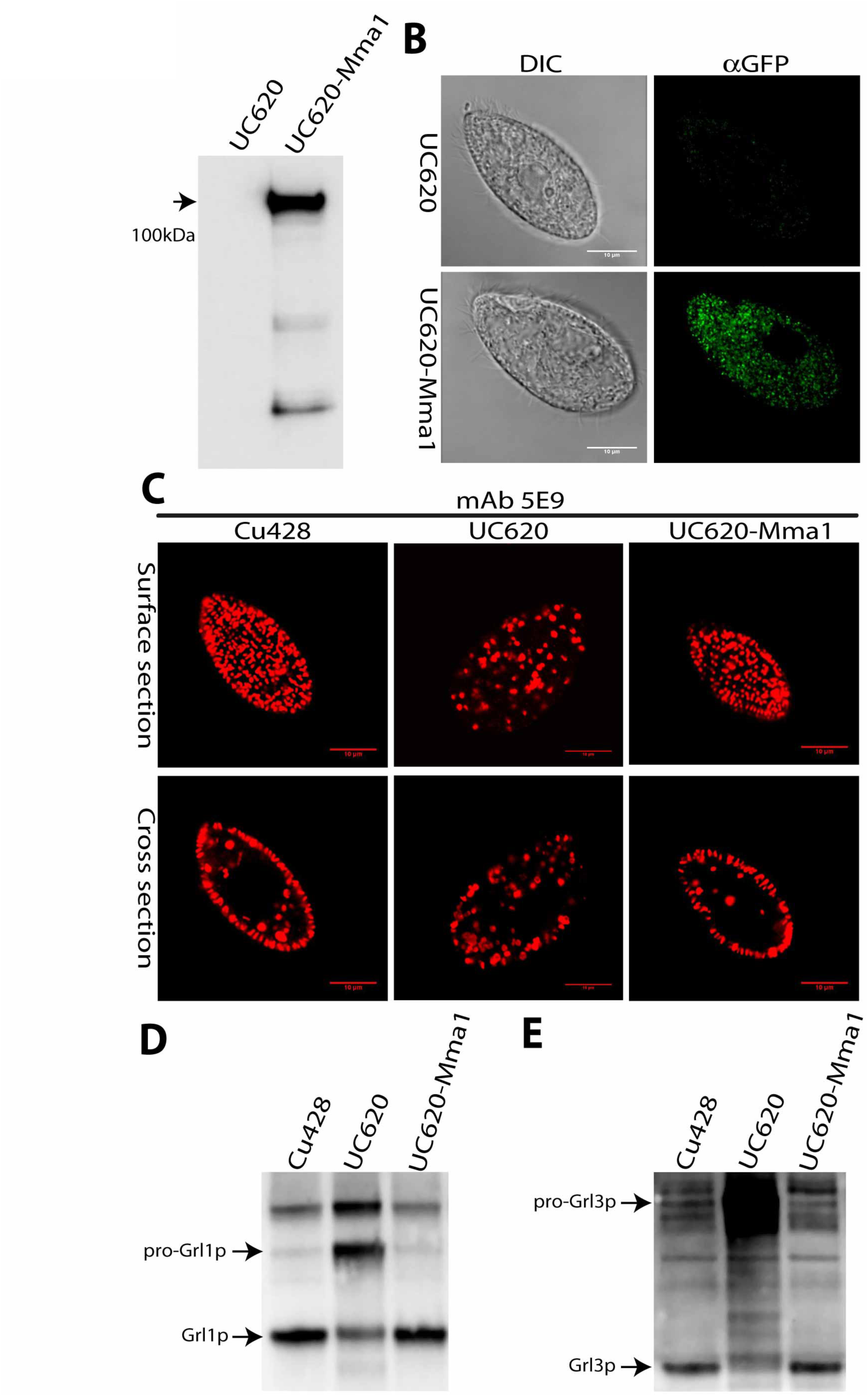
Expression of *MMA1*-CFP rescues the UC620 mutant. (A) Detergent lysates of UC620, or UC620 expressing *MMA1*-CFP, were immunoprecipitated with anti-GFP Ab, and the precipitates separated by SDS-PAGE and immunoblotted with anti-GFP mAb. Cells expressing *MMA1*-CFP show a band of the expected size for the fusion protein (arrow). (B-C) After induction of *MMA1*-CFP with CdCl_2_, cells were fixed, permeabilized, and immunolabeled with rabbit anti-GFP (B) or mAb 5E9 (C), followed by 2° fluorophore-coupled Abs. The scale bars represent 10μm. (B) Mma1p-CFP localizes to multiple cytoplasmic puncta. (C) The expression of the fusion protein in UC620 cells restores the wildtype pattern of docked mucocysts. (D-E) *MMA1*-CFP expression was induced for 2h in growing cell cultures, and WT and UC620 cultures were in parallel treated with CdCl_2_. Whole cell lysates were resolved by 4-20% SDS-PAGE, and transfers were immunoblotted with antibodies against Grl1p (D) or Grl3p (E). The expression of *MMA1*-CFP in UC620 rescues the processing defect seen for both Grl proteins in UC620. Each lane represents 10^3^ cell equivalents.

## Discussion

*Tetrahymena thermophila* possesses many characteristics that make it an attractive model organism for investigating cell biological questions(72). One particularly promising approach to dissect a variety of pathways has been forward genetics following chemical mutagenesis. The cells are diploid, but can be rapidly brought to homozygosity via a specialized mating, thereby facilitating screening for recessive mutations(73). More generally, starved cultures of compatible mating types will undergo highly synchronous mating, and the cells are large enough to be readily isolated as individual pairs, which facilitates classical genetic analysis(53).

Using these and related tools, more than 20 secretion mutants were previously generated by nitrosoguanidine mutagenesis and analyzed to various degrees(41-43). All of them fail to release mucocyst contents upon stimulation, and the underlying cell biological defects range from a block in mucocyst synthesis to defects in exocytosis per se. However, none of the genetic lesions has ever been identified, thus limiting the insights gained from this collection. Similarly, the literature contains detailed analysis of *Tetrahymena* mutants with defects in features such as cortical pattern formation, cell size, lysosomal enzyme release, cell surface antigen expression, food vacuole formation, motility, and rDNA maturation(74-79), but in only a single recent case has the responsible gene been identified(45). That work, by Pearson and colleagues, used a whole genome sequencing approach similar to that which we independently pursued and report in this paper. Our results therefore suggest that mutagenesis linked with whole genome sequencing should be considered a highly accessible approach to link defined pathways with their underlying genes in this organism. This is relevant both for revisiting previously characterized mutants, as well as for developing new genetic screens. Recently, next generation sequencing applied to the ciliate *Paramecium tetraurelia* has illuminated a set of mutants first characterized many decades ago(80).

Our sequencing strategy involved first outcrossing UC620 to generate heterozygous F1 clones, and then using an F1xF1 cross to regenerate homozygous mutant progeny. In principle the second step could have been simplified by crossing the F1 lines with a so-called star (*) strain, employing a strategy called uniparental cytogamy(73). Conjugation in *Tetrahymena* involves the reciprocal exchange of haploid meiotic products between the pairing cells. A cell that conjugates with a * cell fails to receive a viable haploid pro-nucleus, and this promotes the endoreduplication of its own haploid genome. As a result, such * crosses can produce whole-cell homozygotes, which would be ideal for screening and sequencing(73). Star crosses are also known to produce additional, poorly characterized outcomes at low frequency, but this has not detracted from their usefulness for many applications. However, in early experiments we found that a significant fraction of the progeny of the uniparental cytogamy crosses, isolated as individual pairs, showed drug-resistance phenotypes inconsistent with simple endoreduplication, and we therefore employed the somewhat more laborious F1 x F1 strategy.

One key element in quickly identifying the causative SNV in UC620 was our prior physical mapping. Such mapping is relatively straightforward in *Tetrahymena* due to the availability of a panel of so-called ‘nullisomic’ strains, each bearing a homozygous deletion of an entire micronuclear chromosome, and available from the Tetrahymena Stock Center(81). By crossing a strain bearing a recessive mutation with the panel of nullisomics, one can quickly identify the micronuclear chromosome on which the mutation resides. By this approach, the UC620 mutation was mapped to chromosome 1(43). By using an additional panel of available strains with partial chromosomal deletions, this approach may be extended to map mutations to chromosome arms. However, we failed to obtain viable progeny using some of the partial deletion strains on chromosome 1. In the future, it might be very valuable to generate a larger panel of partial chromosomal deletions to cover the *Tetrahymena* genome. If a fine scale panel were available, the mapping of a mutation might be sufficient to then directly identify a small number of candidate genes by sequencing. This approach would bypass the multiple crosses, mating type testing, and progeny screening that we employed to identify *MMA1*.

The mutation identified in the UC620 strain was predicted to change the *MMA1* stop codon, thereby potentially producing a longer polypeptide product. We found unambiguous *MMA1* homologs in the recently sequenced genomes of three other Tetrahymena species, and these alignments helped to confirm the stop codon assignment. The stop codon mutation may result in production of an unstable product, since we found that the UC620 mutation was phenocopied by complete deletion of the *MMA1* gene. *Δmma1* cells, like UC620, showed defects in biochemical and morphological maturation of mucocysts, which were conditional with respect to growth conditions. Moreover, the defects in UC620 were fully rescued by expression of a wildtype copy of *MMA1* tagged with CFP.

Judging by transcript abundance, *MMA1* is expressed at a very low level. Consistent with this, we could not detect Mma1p tagged with 3xGFP, when expressed under its endogenous promoter, and had to induce over-expression to visualize the protein in live cells. Analysis of those cells suggested that Mma1p is a cytosolic protein that partially associated with membranes, which may be small vesicles. This assignment as a cytosolic protein is also consistent with *in silico* predictions, namely the apparent absence of an endoplasmic reticulum translocation signal sequence or of any likely transmembrane domains. The precise role of Mma1p is unknown. Mucocyst biogenesis depends in part on endolysosomal trafficking, as judged by the requirement for a receptor in the VPS10 family and several other proteins associated with trafficking to lysosome-related organelles(58)(unpublished). The small relatively homogeneous Mma1p-GFP-labeled puncta may therefore represent endosomes involved in mucocyst formation.

We have previously noted that the genes encoding all known luminal proteins in mucocysts were co-expressed, together with a receptor responsible for their delivery and the processing enzymes responsible for their maturation(28, 58). Since the expression profile of *MMA1* closely matches that of the *SOR4* receptor, it appears that the phenomenon of co-regulation extends to genes encoding cytosolic machinery in this pathway.

Although a mechanistic understanding of Mma1p action during mucocyst maturation is not yet in hand, our results demonstrate the power of forward genetics in *T. thermophila* to identify lineage-restricted genes that play essential roles in this pathway. It is particularly interesting that *MMA1* has no identifiable homolog in *Paramecium tetraurelia*, another Oligohymenophorean ciliate that makes secretory granules, called trichocysts. Trichocysts, like mucocysts, undergo biochemical and morphological maturation. Significantly, for virtually all known components of *Paramecium* trichocysts and *Tetrahymena* mucocysts, one can readily identify homologous (and likely orthologous) genes in the other organism. This is true for genes encoding the luminal cargo, the processing enzymes, and membrane proteins involved in docking. Thus the absence of an *MMA1* homolog in *Paramecium* is exceptional, and suggests recent emergence of a novel mechanism in an otherwise conserved pathway in the Oligohymenophorean ciliates.

## Acknowledgements

We gratefully acknowledge valuable advice from Eric Cole (St. Olaf College, MN) on the interpretation of uniparental cytogamy crosses, and help from Eileen Hamilton and Ed Orias (UC Santa Barbara, California) on genetic crosses and on mapping Macronuclear sequences to Micronuclear chromosomes. Wei Miao (Chinese Academy of Sciences, Wuhan, China) and Shelby Bidwell and Robert Coyne (J. Craig Venter Institute, Rockville, MD) provided valuable help in annotating *MMA1* and homologous genes in *T. thermophila* and other Tetrahymenids; Doug Chalker (Washington Univ., St. Louis) shared tagging vectors, Jacek Gaertig (Univ. Georgia, Athens) shared anti-tubulin antibodies, Marlo Nelsen and Joseph Frankel (U. Iowa, Iowa City) shared mAbs 4D11 and 5E9, and Kazufumi Mochizuki (IMBA, Vienna, Austria) shared the NEO4 gene disruption construct. Vytas Binokas and Christine Labno (Univ. Chicago Light Microscopy Core Facility) provided expert help, and members of APT’s laboratory benefited from discussion with Joseph Briguglio, Harsimran Kaur, and Daniela Sparvoli. Work in APT’s laboratory was supported by NIH 1R03MH094953 and by NSF MCB-1051985; work in JKP’s laboratory was supported by the Howard Hughes Medical Foundation and NIH 1RO1ES025009.

---

**Fig S1.**
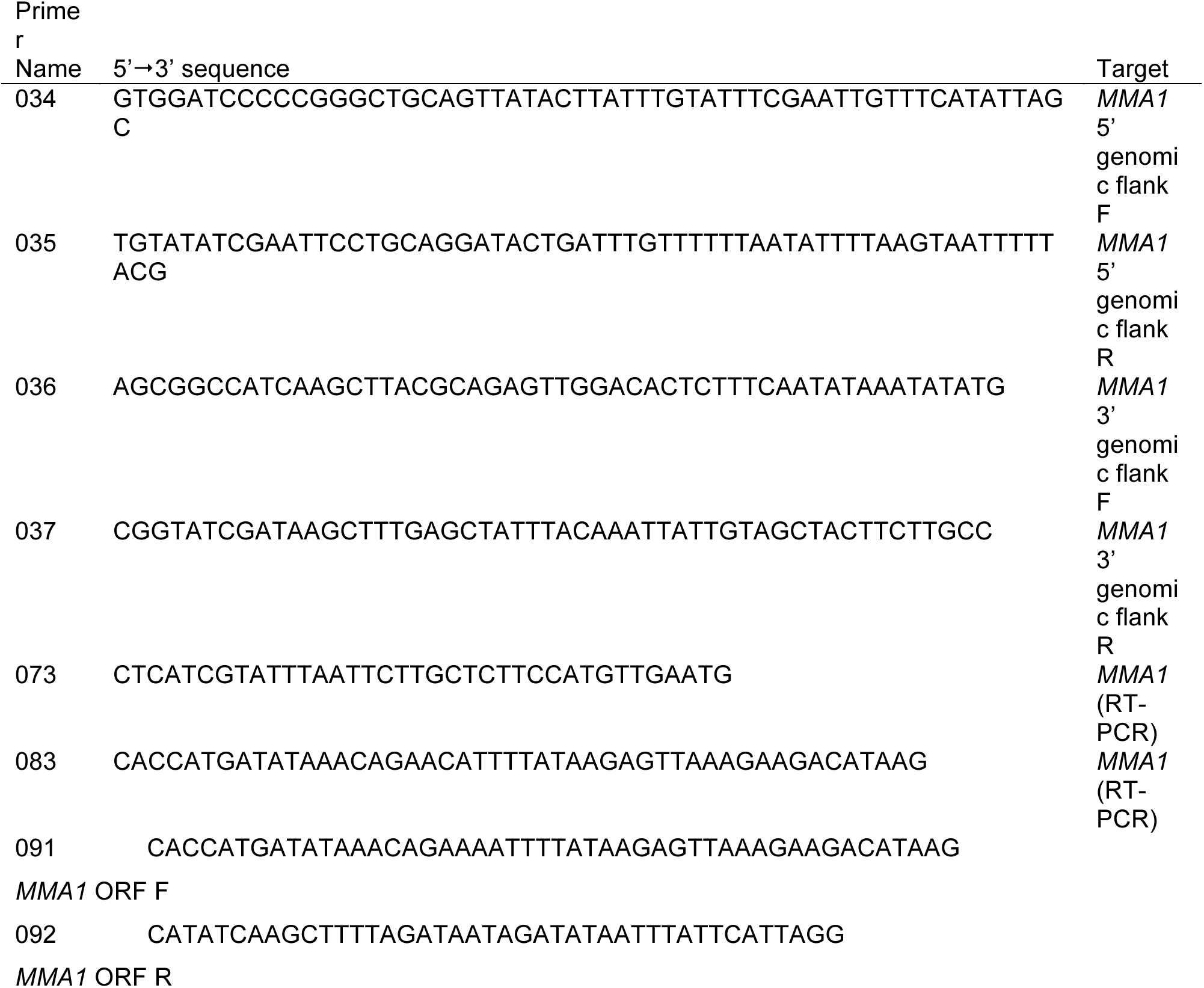

**Figure S2.**
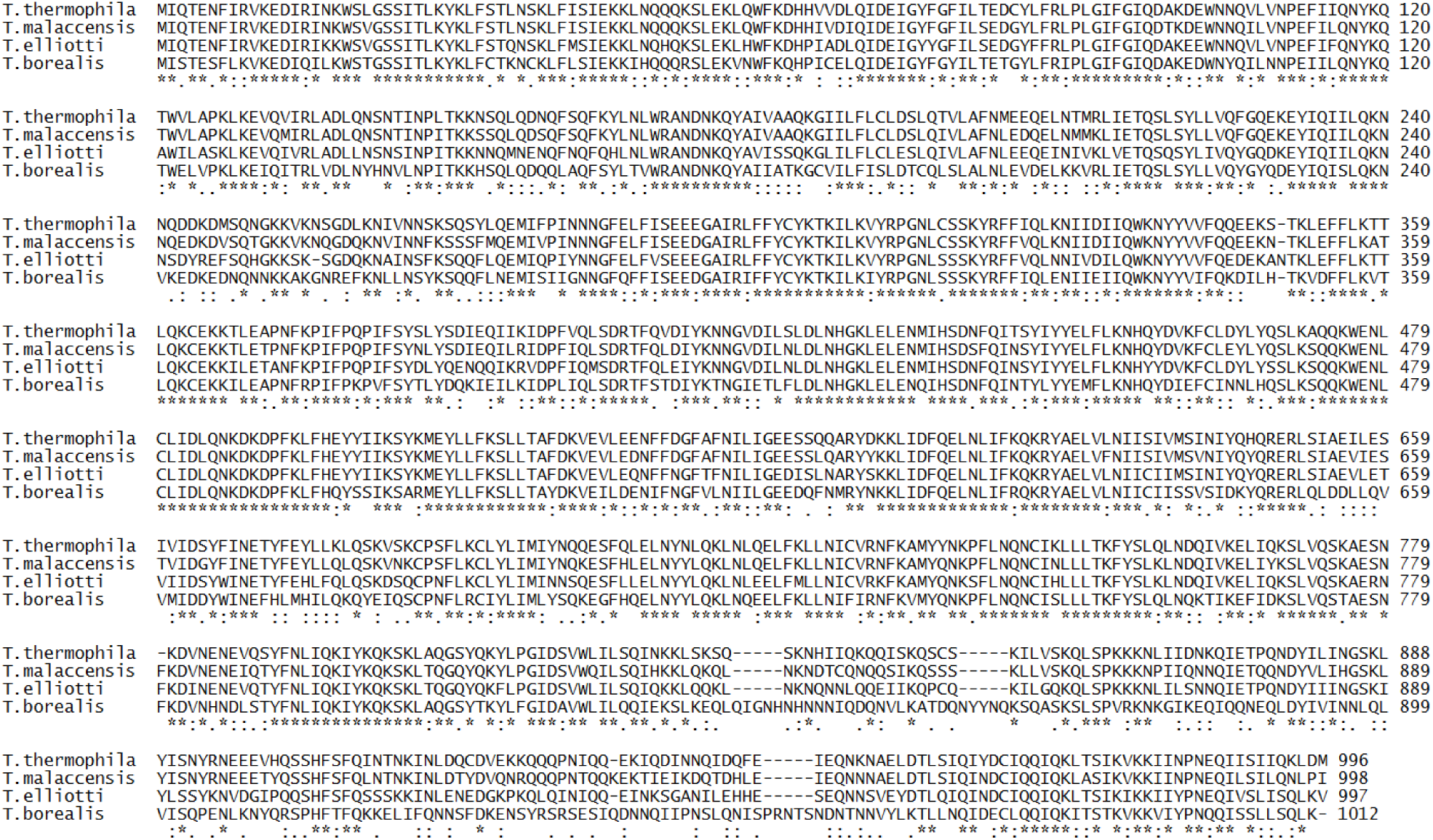
Primary sequence alignment of *MMA1* homologs of four Tetrahymena species (*T. thermophila*, *T. malaccensis*, *T. ellioti*, and *T. borealis*). The GenBank^™^ Accession Numbers for *T. thermophila, T. malaccensis*, *T. ellioti*, and *T. borealis* GM are TTHERM_00566910, EIA_15322.1, EI7_00003.1 and EI9_14649.1, respectively.

